# Roles matter: Graduate student perceptions of active learning in the STEM courses they take and those they teach

**DOI:** 10.1101/502518

**Authors:** Lorelei Patrick, Leigh Anne Howell, E. William Wischusen

## Abstract

Despite many calls to reform undergraduate science, technology, engineering, and math (STEM) education to incorporate active learning into classes, there has been little attention paid to graduate level classrooms or courses taught by graduate students. Here, we set out to understand if and how STEM graduate students’ perceptions of active learning change in the classes they take versus those they teach by administering surveys to STEM graduate students at our institution. We found that graduate students had taken relatively few graduate level classes using active learning and they felt that more time should devoted to active learning in the courses they were taking. Teaching assistants felt that they were devoting the right amount of class time to active learning in the classes they taught. Graduate students also felt that they were using teaching methods in the classes they taught that were different from those they thought should be used when teaching undergraduates and were different from how they preferred to learn when taking classes. These findings suggest that graduate students’ perceptions of teaching and learning changed based on their role in the classroom, which have implications for graduate level course work and professional development programs.

## Introduction

> “Active learning is an instructional method in which students become engaged participants in the classroom. Students are responsible for their own learning through the use of in-class: written exercises, games, problem sets, clickers, debates, class discussions, etc.”
>
> — -Miller and Metz (2014)

Active learning (AL) techniques encompass a variety of teaching practices in which students actively engage with the course content and each other to construct their own knowledge. AL increases student engagement, learning, and course grades, and reduces failure rates (e.g., 1–5); these findings are so ubiquitous (but see 6), that researchers have shifted their focus from proving the benefits of active learning to determining which techniques are most useful for which students in which contexts (7). Despite the preponderance of evidence, AL is not yet used in all classrooms, particularly at the college level (8,9) and much work has sought to identify and lower barriers to implementing AL and other evidence-based teaching practices for faculty teaching undergraduate courses (10–14). Commonly identified barriers for faculty include class size, insufficient class time, and insufficient time to prepare materials (11,13,15).

Faculty and student perceptions of and attitudes toward AL have also been identified as potential barriers to AL adoption (14–20). Unsurprisingly, if faculty don’t perceive that AL is useful or beneficial, they report not using AL in their classrooms; a survey of faculty across disciplines found that attitudes toward AL was the most important predictor AL adoption (18). Student perceptions of AL are also important and can influence whether faculty implement such practices in their courses; concern that students will avoid their courses due to more “difficult” or involved coursework or students simply not liking the technique have prevented some faculty from implementing pre-prepared AL modules (14). Students themselves may perceive that they learn and retain information better through AL, however this doesn’t necessarily mean that they like or fully engage in classes that implement AL (15,17,19,20). Much of this resistance seems to stem from the desire to passively acquire information from an expert – the instructor – which is perceived to require less effort than actively constructing their own knowledge and dealing with the uncertainty of having the “right” answer, in spite of the fact that these are features not only of AL, but also of the scientific process in general (17,19). This suggests that students often fail to realize or appreciate that the thinking skills associated with AL are important for their professional development. This also suggests that students who are using the scientific process on a daily basis – like STEM graduate students – may perceive AL more favorably in the classes they take and those they teach.

Graduate students, particularly graduate teaching assistants (TAs), have a unique position in academia, especially in STEM fields, because they are transitioning from the single role of being strictly students taking courses (acquiring knowledge) to multiple roles, by adding the additional roles of discipline experts through both discovering (researcher) and disseminating (teacher) knowledge. As such, they still must complete coursework, conduct novel research, and in many cases teach at the undergraduate level. Many graduate students in STEM disciplines teach; for example, in biology, 70-90% of laboratory courses in large research universities are taught by TAs (21,22) and TAs primarily teach introductory level lab courses (23). Previous work suggests that graduate students, particularly TAs, can feel out of place – that they are neither “fish nor fowl” – while navigating their roles as student, teacher, and researcher (24). Because of this unique role, graduate students are in an ideal position to inform our understanding of how teaching practices −and perceptions of these practices− develop, particularly in answering the perennial question of whether teachers teach the way they were taught (25). To date, however, relatively little work has examined how graduate students perceive and use AL, or if they are even aware of AL techniques.

Mirroring the findings for faculty (discussed above), TA buy-in and their perception of AL are crucial for proper implementation of evidence-based teaching practices in undergraduate classrooms. Graduate students and TAs are aware of active learning and other evidence-based teaching practices, but not all have bought-in or currently use AL when they teach (26). Course style, previous experiences as a student, previous teaching experiences, and professional development sessions influence TA perceptions of how they should teach (lecturing vs. questioning students) and their role in the classroom (content expert vs. guide 27,28). TAs differed in their opinion of PBL – a type of AL – in a chemistry lab course, ranging from feeling that it is beneficial to their students to feeling that they were being asked to teach in a manner that they perceived as a more difficult and time-consuming (29). In another case, TAs generally enjoyed teaching an inquiry-based lab course, perceived that their own research skills improved, and that their students learned better (30). There have also been multiple studies detailing training programs, workshops, and other professional development activities designed to help TAs become more effective teachers and introducing them to evidence-based, student-centered teaching practices in an effort to better prepare future faculty and improve undergraduate STEM education (31–34); these training programs were successful in improving undergraduate success or influencing TA attitudes toward and confidence in teaching. These studies suggest that more research is needed to fully explore TA perceptions of AL and how these perceptions influence the teaching techniques TAs use.

Recently we explored undergraduate and faculty perceptions of active learning in the courses they take or teach in engineering (in prep) and science (15) at our university. We found that faculty and undergraduates generally perceived that AL was useful and that more class time should be devoted to AL than was currently occurring in their courses. We now focus on graduate student perceptions of active learning in both the classes they take and the classes they teach in order to address the following question: Do the perceptions of active learning by graduate students differ depending on their role (student vs. teacher)? We have divided this over-arching question into the following sub-questions: 1) Do graduate students perceive that they have experienced or used active learning as students and teachers? Do they think it’s effective? 2) Does the value of active learning by graduate students differ depending on their role (student vs. teacher) in terms of class time devoted to active learning? 3) Do the perceptions of active learning by graduate students differ depending on their role (student vs. teacher) in terms of teaching methods? To answer these questions, we surveyed graduate students’ perceptions of active learning in the courses they’ve taken and the courses they’ve taught. The insights we gain from this study should be used to inform the advancement of professional development for TA and non-TA graduate students.

## Methods

### Institution

This study was conducted at a large research-focused university (basic Carnegie classification: Doctoral Universities: Highest Research Activity) in the southeastern United States. We focused on two colleges within the university: the College of Engineering (CoE) and the College of Science (CoS). The CoE consists of seven departments and the CoS of five departments (Table 1).

**Table 1.**
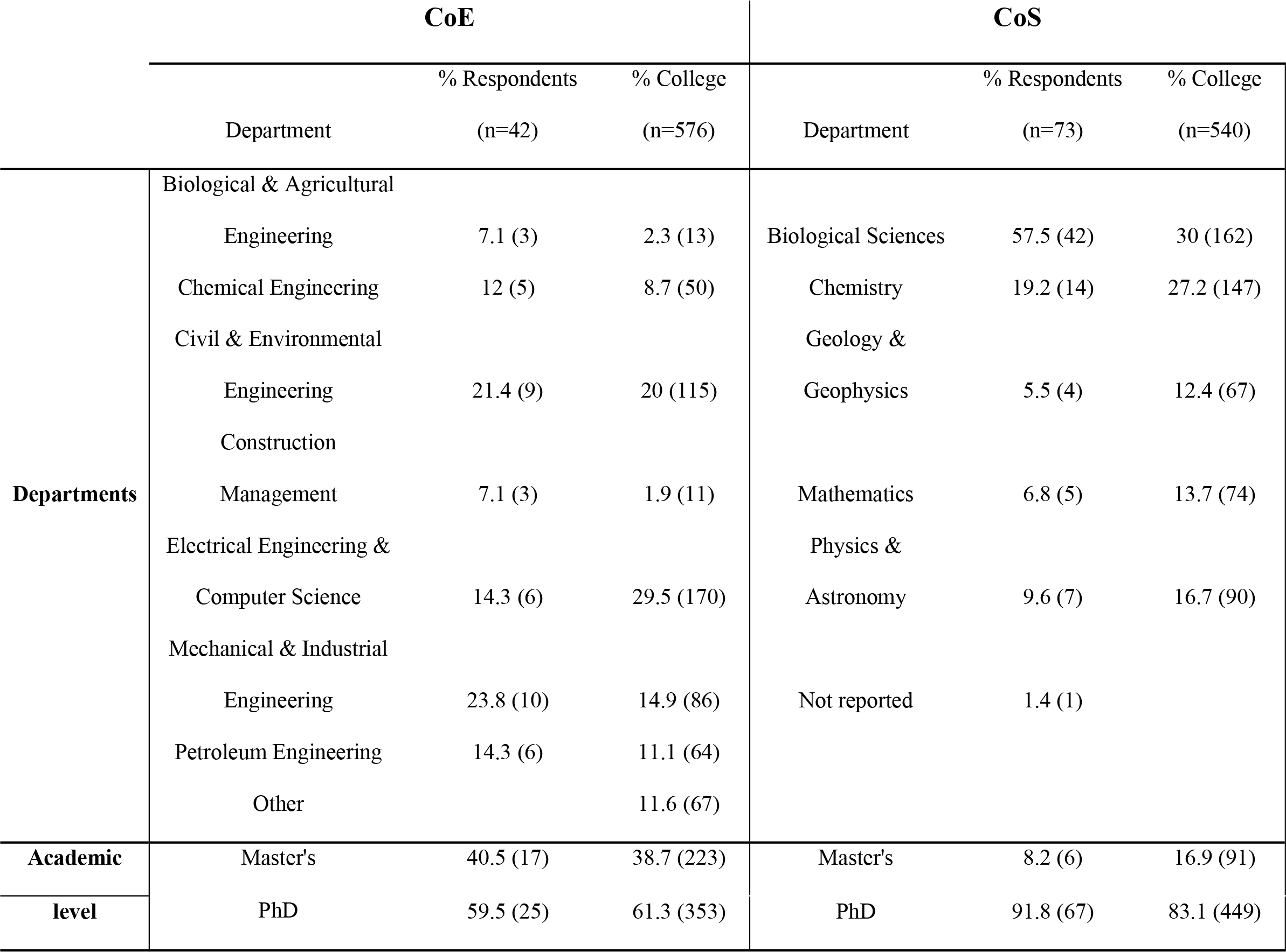
Demographic makeup of the respondents to the survey and of the Colleges at the time the survey was distributed.

Percent of respondents (and number of respondents) are presented. “Other” respondents were from interdepartmental degree programs within the College of Engineering. “Not reported” indicates respondents who declined to give this information.

All departments in both Colleges provide teaching assistantships to graduate students, but in some departments there are more graduate students than assistantships, so that not all graduate students will teach as part of their graduate education. Biological Sciences (CoS), Chemistry (CoS), and Chemical Engineering (CoE) are the only departments that require all graduate students to teach for at least one (Biological Sciences) or two (Chemistry and Chemical Engineering) semesters before graduation. Mathematics (CoS) requires all graduate students to take a two-course sequence on math pedagogy and Biological Sciences (CoS) requires all graduate students lacking prior teaching experience to attend a 1-credit course on science pedagogy. All departments encourage, but do not require, graduate student participation in orientations and workshops organized by the Graduate School. In most cases, TAs are responsible for running lab or recitation sections, but occasionally they may be the instructor of record and solely responsible for a lecture or laboratory course.

### Study Participants

The target group consisted of all graduate students within the CoE and CoS. Graduate students in the CoE were emailed the survey using the College’s graduate student listserv during the Spring 2015 semester. Surveys were emailed directly to graduate students in the CoS during the Fall 2014 semester using lists assembled from department websites or provided by individual departments. Surveys were available for completion for two weeks after they were initially sent; follow-up reminders were sent after one week and one day before the survey closed.

### Survey instruments

We modeled our survey instrument after those used by Miller and Metz (13) and modified for use at our institution by Patrick et al. (15). Since we were interested in graduate student perceptions from both their student and teaching perspectives, we combined the instrument for students – asking about AL in the courses they take – with the instrument for faculty – asking about AL in the courses they teach (13,15). To help ensure that all survey participants approached thinking about AL from the same perspective we provided Miller and Metz’s (13) definition of active learning at the beginning of each survey section:

> “Active learning is an instructional method in which students become engaged participants in the classroom. Students are responsible for their own learning through the use of in-class: written exercises, games, problem sets, clickers, debates, class discussions, etc.”

The student role section was presented immediately after the consent form, followed by the teaching role section. In the student section, respondents were asked how long they had been in graduate school, how many classes they had taken, how many courses had used active learning, to rank teaching practices based on how well they learned when the practice was used, and how much class time was currently and should be devoted to AL in graduate level courses. Using class time as a proxy for acceptance of AL may be a somewhat problematic metric because one can imagine a scenario in which someone might be an advocate for AL but think that only a small, but presumably a non-zero, proportion of class time should be devoted to it. It is harder to imagine a scenario in which someone who does not buy in to AL would devote a large proportion of class time to it unless required to do so. Despite this caveat, we use class time as a cautious estimate of AL buy-in.

The teaching section asked similar questions but from the perspective of an instructor; that is, how respondents thought undergraduates should be taught and how the graduate students had taught or were currently teaching undergraduates, how many semesters they had taught, and how many different courses they had taught. We did not ask detailed demographic information to protect each respondent’s identity. The instrument is available in the online Supplemental Materials 1. The original survey was designed for post-baccalaureate students pursuing advanced degrees (dental students) and their professors (13), however we did not independently validate the instrument. This project was conducted with approval of our institution’s IRB, project #E9078.

### Statistical analyses

Data were downloaded from Qualtrics to Excel and partial responses were removed. We used the Wilcoxon Signed-Rank test to compare differences in paired data between respondents’ student role and teacher role perceptions. All analyses were carried out in R (35). Figures 1 and 2 were made in R; Figure 3 was made using RAWGraphs (36).

**Figure 1.**
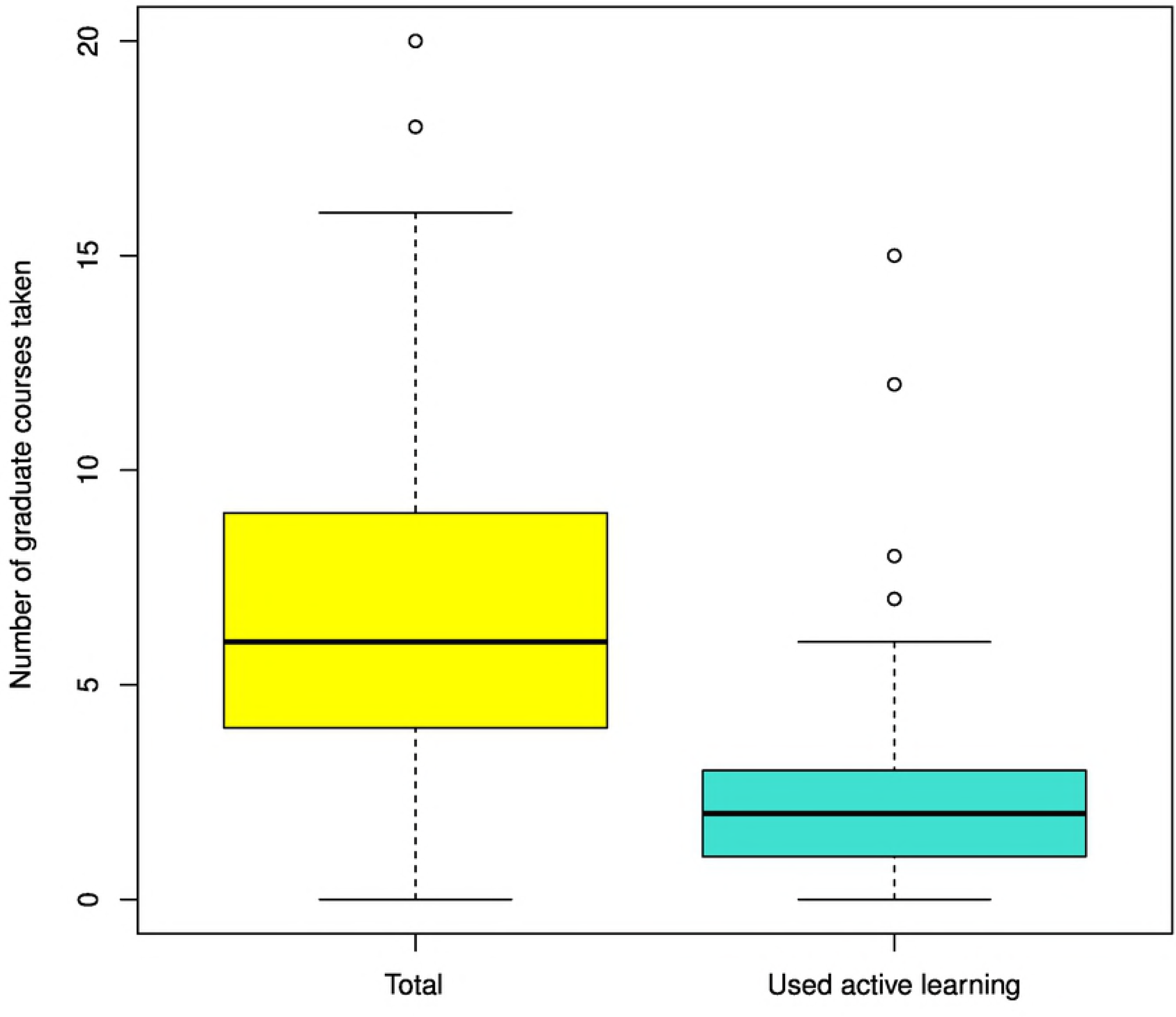
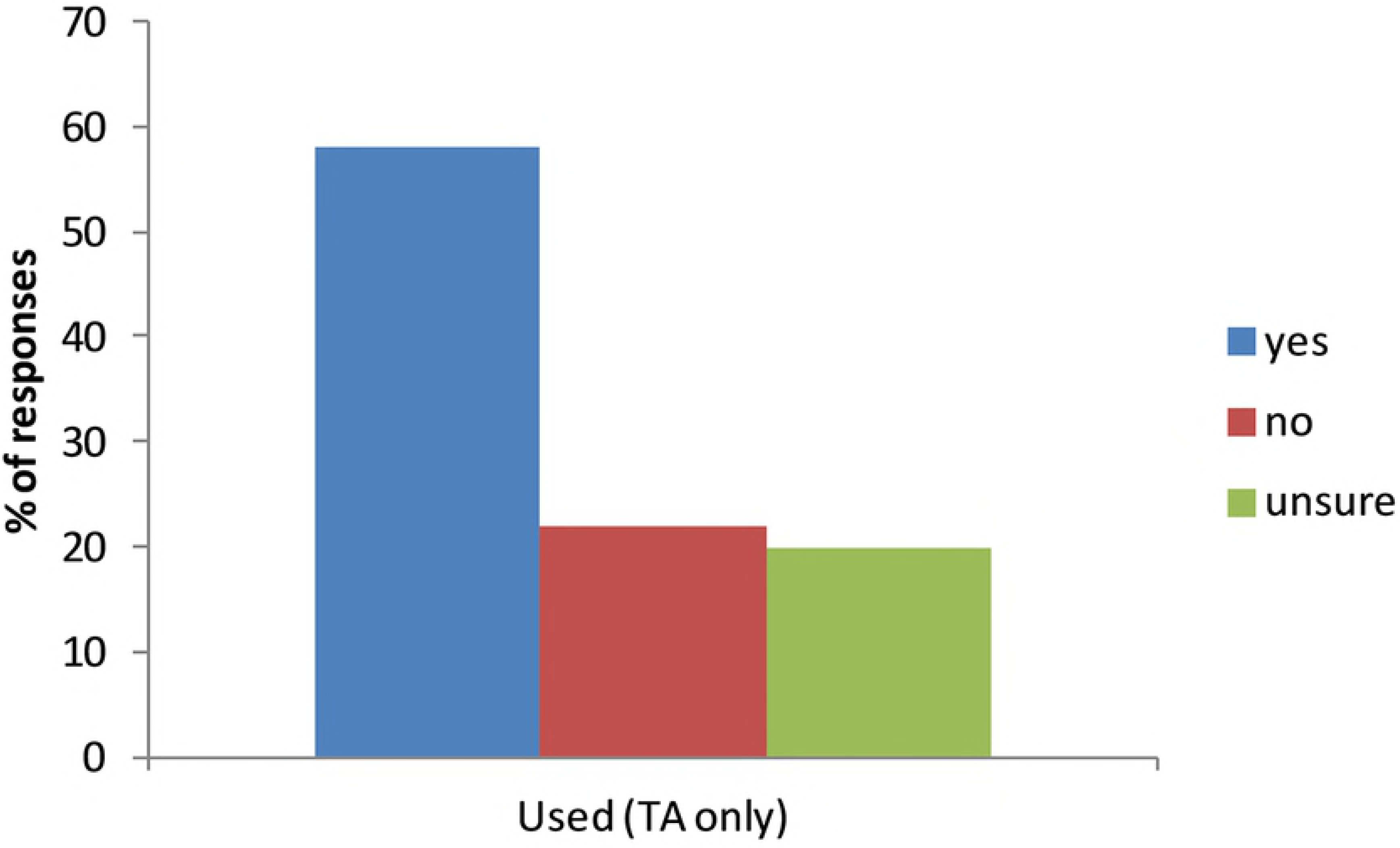
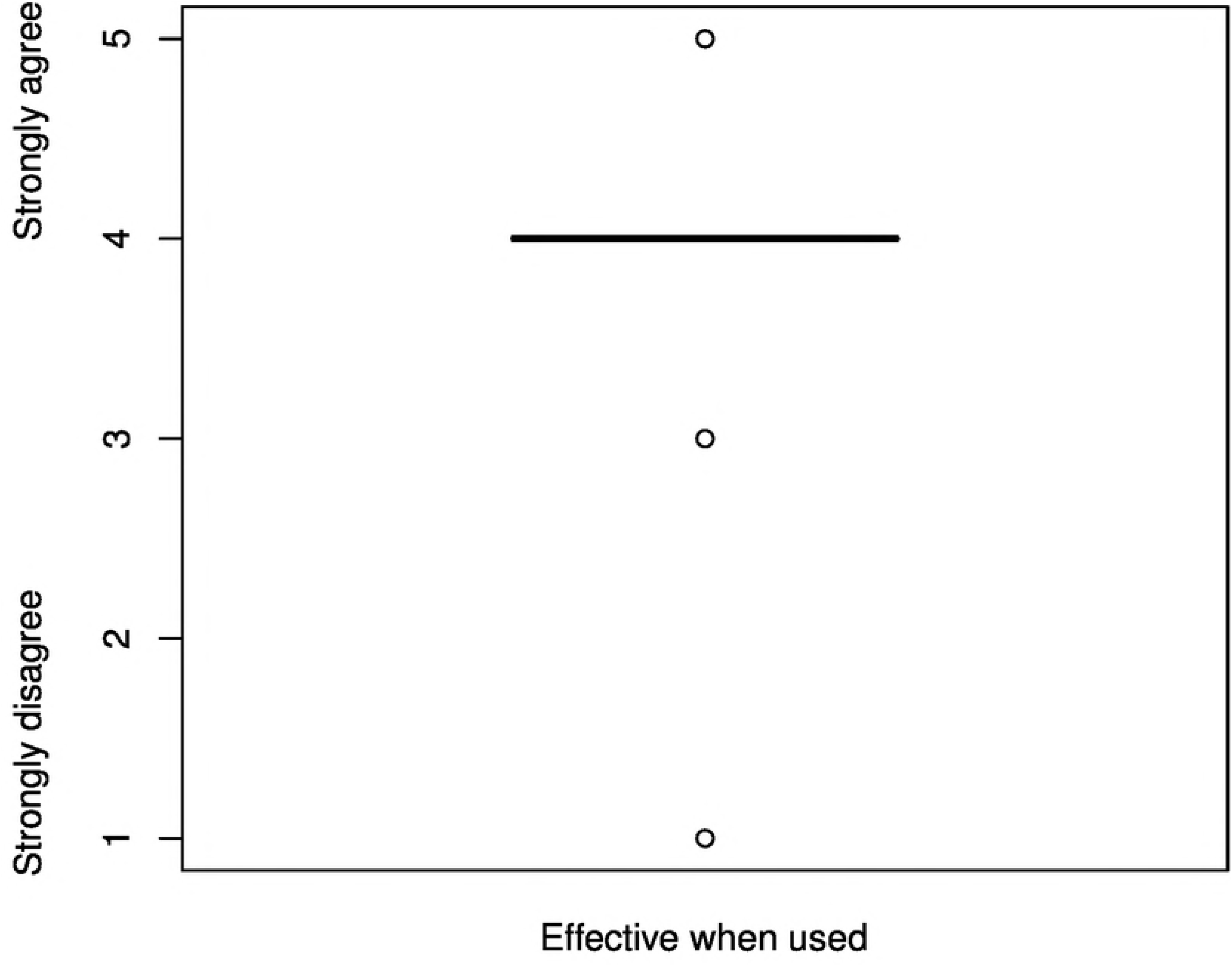
A) Total number of graduate level courses taken at our institution and number of courses that used active learning teaching practices (N=114). In box plots, the thick horizontal bar indicates the median, the shaded box surrounding the median indicates the interquartile range between the first and third quartiles, and the whiskers depict 1.5 times the interquartile range. B) Percent of teaching assistants (TAs) who used active learning (N=81). C) Box plot of TA perceptions of the effectiveness of active learning when used. (N=81).

**Figure 2.**
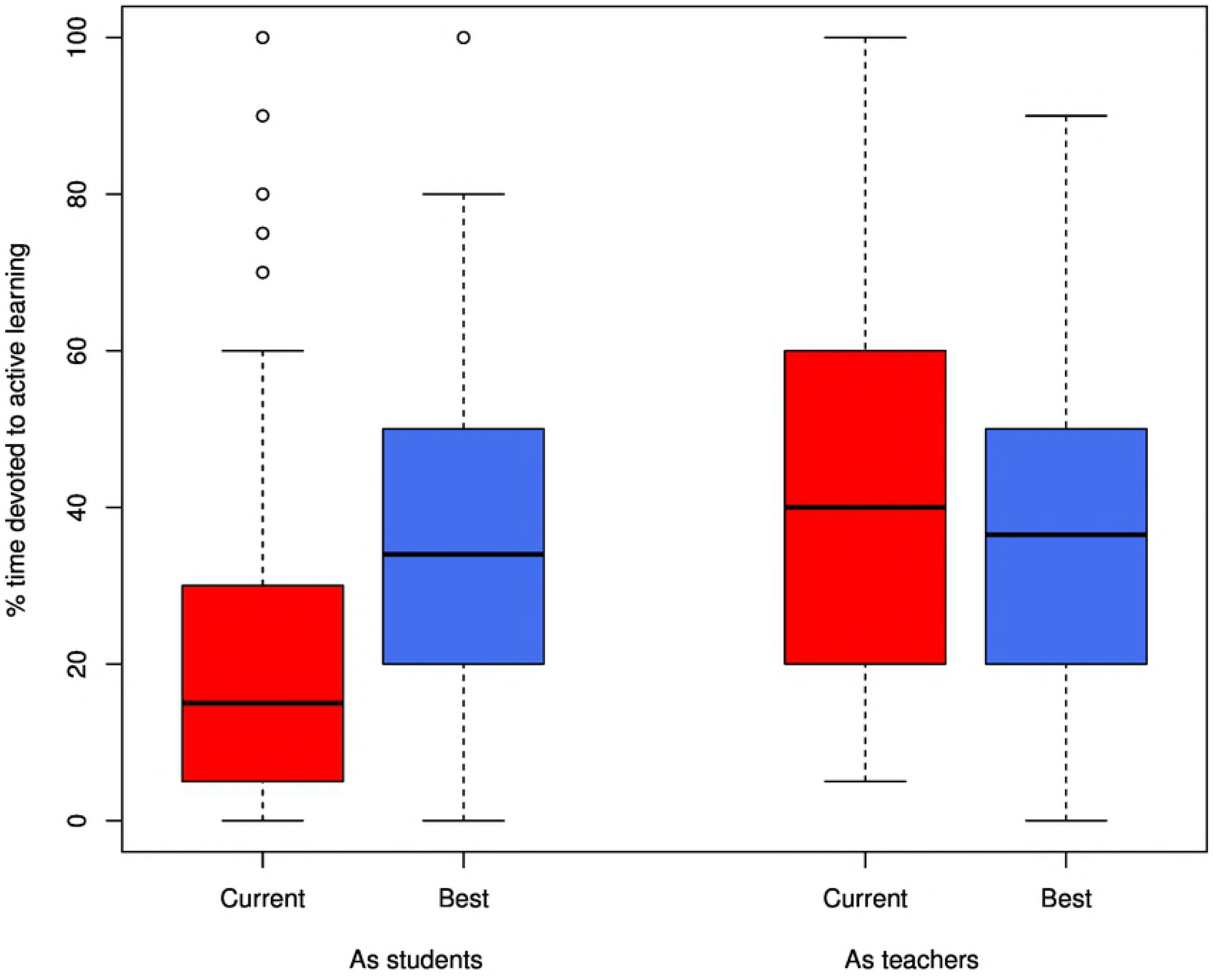
Box plot of graduate student perceptions of the amount of class time currently devoted to active learning (“Current”) and how much time should be devoted (“Best”) to active learning in the classes they take (“As students”) and the classes they teach (“As teachers”). Wilcoxen Signed-Rank tests indicate there was a significant difference between the “Current” and “Best” “As student” boxes (the two boxes on the left; p < 0.001) and the “Current” “As students” and “Current” “As teachers” boxes (the red boxes; p=0.002). In box plots, the thick horizontal bar indicates the median, the shaded box surrounding the median indicates the interquartile range between the first and third quartiles, and the whiskers depict 1.5 times the interquartile range.

**Figure 3.**
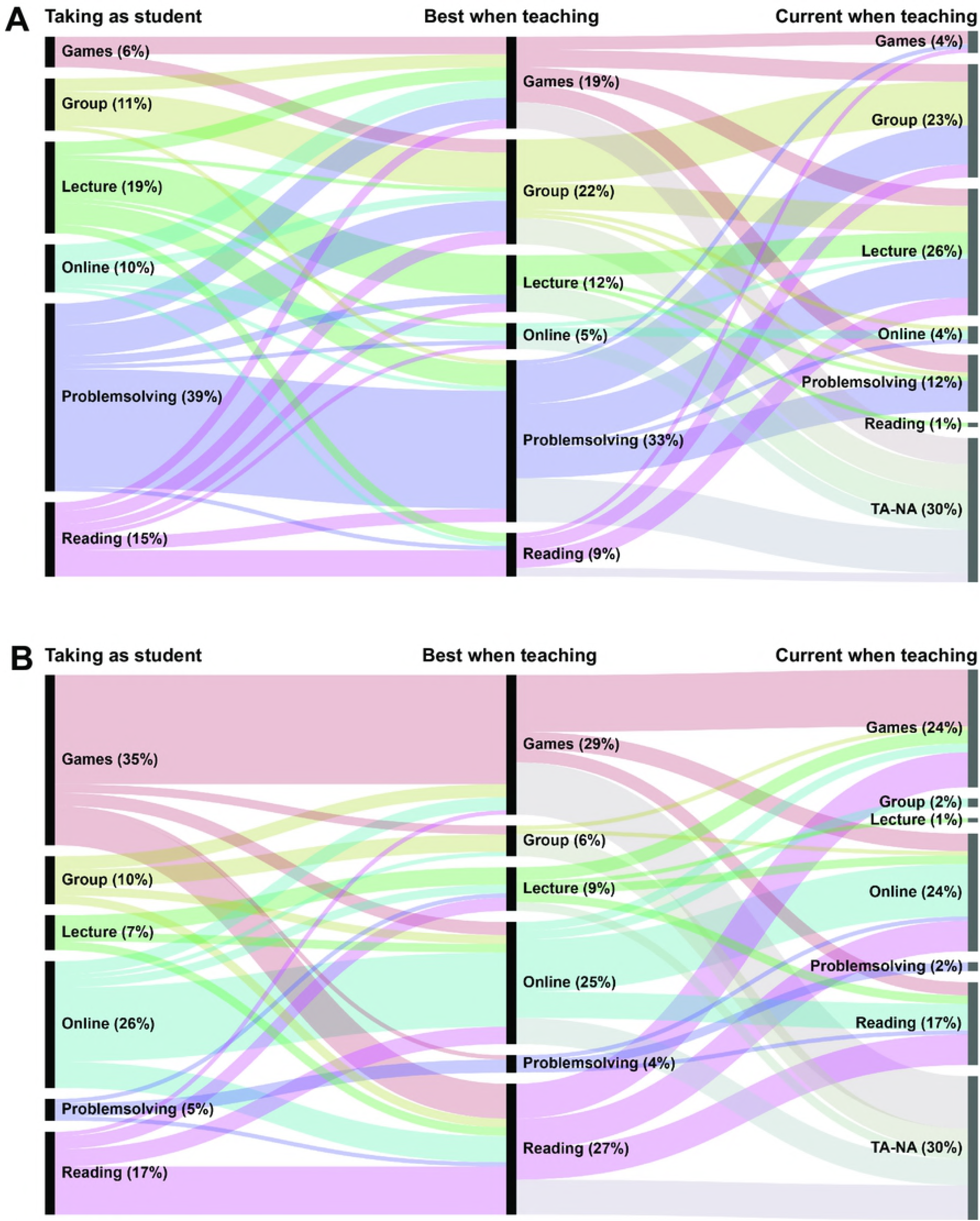
Alluvial plots of A) the *top* (ranked 1st) teaching practices for graduate students in different roles and B) the *bottom* (ranked 6th) teaching practices for graduate students in the same roles. The vertical bars are the teaching practices; the vertical height of each bar is proportional to the number of respondents who chose that teaching method. In both A) and B), the first column is the preferred learning method *when taking classes*, the middle column is what graduate students consider to be the *best way to teach undergraduates*, and the last column are the teaching methods *TAs were currently using*. The “alluvia” or lines between columns indicate how graduate student perceptions of teaching practices differed or remained the same between roles and are proportional to number of respondents. TA-NA= respondents who were not or had never been teaching assistants.

## Results

### Number of participants

A total of 42 (7%) graduate students from the College of Engineering (CoE) and 73 (13.5%) graduate students from the College of Science (CoS) responded to the survey. The majors and degree programs of the respondents were fairly representative of the overall composition of graduate students in the CoE (Table 1). Student responses from Biological Sciences were overrepresented compared to the overall composition of the CoS (Table 1) but otherwise the departments and degree programs were reasonably well represented (Table 1). Only 17 (40%) of the respondents from the CoE were currently TAs or had served as TAs in the past whereas 64 (90%) of the respondents from the CoS were or had been TAs.

### Do graduate students perceive that they have experienced or used active learning? If so, did they think it was effective?

Graduate students had taken a median of six (range: 0-20) graduate level courses at our institution (Figure 1a). Relatively few of these graduate level courses used AL (Figure 1a; median=2; range= 0-15); 26 graduate students (23%) reported that none of their graduate courses used AL. Of the graduate students who were or had been TAs at our institution, 58% reported that they had used AL in their classrooms (Figure 1b). Graduate students agreed that AL was effective when they used the techniques as an instructor (Figure 1c).

### Do the perceptions of active learning by graduate students differ depending on their role (student vs. teacher) in terms of class time devoted to active learning?

We compared graduate student perceptions of the amount of class time devoted to AL in the classes they take (“As students” panel of Figure 2) and the classes they teach (“As teachers” panel of Figure 2). They were asked to estimate the amount of time that was currently devoted to AL in these classes (“Current”) and the amount of time they thought should be devoted to AL (“Best”) in Figure 2. As students, graduate students reported that significantly less class time was being devoted to AL (“Current”) in the classes they had taken compared to how much time they thought *should* be devoted to AL (“Best”; “As students” panel of Figure 2; p < 0.001). As teachers, there were no significant differences between how much time was currently or should be devoted to AL in the classes taught by graduate students (“As teachers” panel of Figure 2). Graduate students reported that significantly more AL was occurring in the classes they were teaching than in the classes they were taking (“Current” bars in the “As students” vs. “As teachers” panels, Figure 2; p= 0.002) but not in their perception of the time that should be devoted to AL.

### Do the perceptions of active learning by graduate students differ depending on their role (student vs. teacher) in terms of teaching methods?

Interesting differences between roles were uncovered when we examined graduate student perceptions of teaching methods (Figure 3). Figure 3A shows the top ranked teaching method for all respondents when taking classes (left column), the best practice when teaching undergraduates (middle column), and how TAs were currently teaching (right column). When graduate students were asked to rank teaching methods by how they learned most effectively, the largest proportion of respondents ranked “Problem solving” as their most preferred learning method (indicated by the size of the vertical bar, 39% of respondents), followed by “Lecture” (19%); “Educational Games or Activities” was the top ranked learning method with the fewest respondents (6%; Figure 3A, left column). When asked to rank best teaching practices for undergraduates to learn, “Problem solving” was again the top ranked method for the largest proportion of respondents (largest vertical bar; 33%), followed by “Group or Collaborative Learning” (22%) and “Educational Games or Activities” (19%; Figure 3A, center column). The alluvia (colored lines indicating proportion of respondents) between the left and middle columns show that many respondents thought that their preferred learning method was also the best teaching method for undergraduates; this relationship is particularly strong for “Problem solving”. However, in other cases such as for “Reading” and “Online learning”, respondents thought that their preferred learning method was not the best practice when teaching undergraduates. Nearly equal proportions of TAs ranked “Lecture” (26%) and “Group or Collaborative Learning” (23%) as their most used teaching methods (Figure 3A, right column) despite thinking that other teaching methods were better (alluvia between middle and right columns, Figure 3A). Problem solving, which was the highest ranked for both preferred method of learning and best teaching practice, was the top-ranked teaching practice for only a third (12%) of current or former TAs (right column, Figure 3A).

Figure 3B shows the proportion of respondents that ranked each teaching method as their lowest or bottom choice. Graduate students do not prefer “Educational Games or Activities” (35%) or “Videos or Online Learning” (26%; tall vertical bars in the left column, Figure 3B). These two methods (29% and 25% respectively), along with “Reading” (27%) were also ranked as the worst methods for teaching undergraduates (middle column, Figure 3B). These three methods, “Educational Games or Activities” (24%), “Videos or Online Learning” (24%), and “Reading” (17%) were also the lowest ranked methods by TAs currently teaching (right column, Figure 3B). Most TAs ranked “Group or Collaborative Learning” (2%), “Lecture” (1%), and “Problem solving” (2%) highly, indicated by the very small vertical bars for these methods in the right column of Figure 3B.

## Discussion

Graduate students at our institution report that active learning is effective and that more time should be devoted to these teaching techniques in the classes they take. Our results also show that graduate students simultaneously have a foot in both student and teacher worlds: their perceptions of AL in the courses they take were similar but not identical to their perceptions of how they should or were currently teaching. To our knowledge, this is the first study to investigate perceptions of AL in graduate students as both students and teachers.

### Do the perceptions of active learning by graduate students differ depending on their role (student vs. teacher)?

In short, yes. However, it is important to reiterate that these were self-reported perceptions of active learning techniques and we cannot independently verify the proportion of graduate level courses actually using AL, amount of time TAs actually devoted to AL, or that respondents consistently used the definition of AL we provided when answering the survey questions. However, recent work has shown that at least some AL is implemented in the majority of STEM middle school, high school, and undergraduate classrooms in Maine (8), university level STEM courses across North America (9), and interviews with graduate students indicate they are aware of AL teaching practices (36), all of which suggest that AL is becoming increasingly prevalent and graduate students know what these teaching practices are. In addition, the majority of CoS faculty and undergraduates at our institution reported using and/or experiencing AL in at least some their courses (15). For these reasons, we are reasonably confident that most graduate students have been exposed to AL at some point in their academic careers and so have some idea (albeit potentially incorrect) of what AL is.

Akiha and co-workers (8) found that when students transition from high school to college, they experienced a marked instructional shift away from the majority of class time being devoted to AL, to lecture dominating their college classroom instruction. Our data suggest that students transitioning from undergraduate degrees to STEM graduate studies may experience another instructional shift to even less AL in their graduate level courses. Previous studies of graduate student perceptions of AL in the classes they take suggest that their attitudes range from neutral through positive. For example, Tune et al. (37) examined graduate student performance and perceptions of a flipped-class versus traditional, lecture-style structure in a physiology course. They found that the graduate students in the AL flipped classroom performed better on exams and perceived they learned more than in a traditional lecture course (37). However they felt strongly that they spent too much time outside of class preparing (37) a perception that has also been reported for undergraduate students (20). Jones et al. (38) employed a different AL technique, problem-based learning (PBL), in an ethics class for biomedical graduate students. Despite initial resistance due to their preference for lecture and unfamiliarity with PBL, anecdotal evidence suggested that students in the course grew to prefer the technique over traditional teacher-centered teaching practices (38). When dental students in a team-taught physiology course were presented a portion of the material in a traditional lecture format and the remaining material presented using several active learning techniques, the students showed a strong preference for active learning, perceived that they learned more, were more actively engaged in the course, and thought that more active learning should be included in graduate level courses (13).

Much time, effort, and resources have been directed at reforming undergraduate education in the past few decades but relatively little has been devoted to education reform at the graduate level. Our results indicate that many (although not all) graduate students have bought in to the premise that AL is beneficial to their learning and want at least some AL in their courses (~36% of the class time). Therefore, more emphasis should be placed on learner-centered graduate-level classrooms. While there is a growing body of literature discussing the incorporation of AL into courses and workshops focused on teaching, relatively few publications outline how graduate level STEM courses not related to teaching have been made more student-centered (i.e., 13,39–41). Such publications would be useful examples to instructors wanting to reform their graduate level courses but unsure of how to start the process.

Although not specifically examining perceptions of active learning, previous work investigating student outcomes and teaching beliefs indicates that there is essentially no difference between undergraduate and graduate TAs (42,43). Combined with our results, these findings imply that changing a person’s role in the classroom may be all that is required to change their perception of teaching and learning. In the current study, when asked to switch from thinking about their student role to their teaching role, graduate student rankings of teaching techniques began to differ. Specifically, TAs were currently teaching differently from the way they preferred to learn and how they thought undergraduates learned best. The premise that we teach the way we prefer to be taught is incorrect, at least for our respondents; a similar finding has previously been reported for faculty members (25). Our findings also suggest that overall, graduate students report more positive perceptions of some AL teaching practices over more “traditional” passive teaching methods.

### Implications and recommendations for graduate and undergraduate education

Since embarking on this project, we have been asked many times “Why should anyone care what graduate students think of active learning?”. Research indicates that AL is beneficial in essentially all contexts (e.g., 3,4,13) and that positive perceptions, positive experiences, and buy-in are important to and predictive of AL implementation by instructors and TAs (e.g., 14,17–19,29,30). Therefore, we argue, and the evidence indicates, that one should care about graduate student opinions for the same reasons that we should care about undergraduate and faculty perceptions: because graduate students are currently in classrooms taking and teaching courses.

If undergraduate education reform progresses as advocates hope, an increasing number of students will enter graduate or professional schools having already been exposed to AL and scientific teaching practices during their undergraduate degrees (8,9). These students will likely expect that their graduate courses will take these practices to the next level and, given student responses to our survey, will also likely be disappointed that this is not always the case. As discussed above, we recommend that faculty reform not just their undergraduate courses, but also their graduate level courses and that these course resources be made widely available. Although faculty and graduate students don’t necessarily teach the way they were taught (25 and our results presented above), modeling desired teaching techniques in graduate courses can certainly only help efforts to reform teaching in the academy. In order to ensure AL occurs in graduate courses, departmental, college, university, and/or funding resources for reforming teaching should not be reserved only for undergraduate classes. Barriers preventing faculty from implementing AL in undergraduate classes, although likely to be similar to those for graduate courses (with the possible exception of class size), should be examined (e.g., 11,13,15) and lowered whenever possible.

Even more importantly, TAs are already in the classroom, teaching undergraduate STEM students in courses with much smaller student-to-teacher ratios than most lecture courses. Therefore what they do in the classroom matters just as much as what faculty do in larger courses. This is especially important because the majority of TAs teach in introductory level courses, often referred to as “gateway” courses because of the barrier they pose to many students (21,23,44). As outlined above, understanding graduate student perceptions toward various teaching strategies is one key to increasing buy-in and developing impactful professional development training in order to increase learning and retention in undergraduate courses and provide TAs with training aligned with their level of exposure to evidence-based teaching practices, which should positively influence future undergraduate education. Our findings strengthen recent calls to provide more pedagogical training to graduate teaching assistants, particularly early in their program (31,34,44). There are several courses, workshops, and frameworks that have been developed and assessed, so departments and colleges looking to start or revise graduate student professional development need not start from scratch (e.g., 23,32,33). Recent research suggests that such training not only enhances undergraduate education in the short- and long-term, but graduate students completing such training and programs graduate on time and are employed at a higher rate than graduate students who don’t participate in such professional development (45,46). In addition to more formal course- or workshop-based TA training, we make the additional recommendation that shorter teaching training sessions on AL be added to the TA meetings that typically accompany graduate teaching assignments and that the coordinators leading these meetings model the types of teaching activities and behaviors TAs can – and should – incorporate into their sections (28). As discussed in more detail below, TAs often have little to no control over *the content* of the courses they teach, however in many cases they do have control over *how they teach* the content in their sections. Therefore we suggest that these in-meeting training sessions demonstrate how TAs can present specific course content in an active manner and facilitate brainstorming sessions in which TAs suggest ways to present other course topics using a variety of active learning teaching methods.

### Study limitations

Possible limitations of the survey instrument in general are discussed at length in a previous publication (15). These limitations include using the word “lecture” to refer to the non-laboratory portion of courses as well as the teaching method, the lack of questions assessing participation in teaching training, the lack of incentives for survey respondents, and the broad “Teaching Methods” categories. Importantly, our survey instrument has not been validated either by us or by the authors of the original instrument (13,15), so our results should be interpreted with caution. In addition, the dual roles that graduate students play could lead to potential limitations of this study and the applicability of our findings to other institutions. First, it is common, but not universal at our institution, for graduate students to teach one or more sections of a course made up of multiple sections. It is much less common for TAs to be the instructor of record, but it does occur occasionally in both Colleges. TAs who are not instructors of record likely have little control over the content covered in the course. Although in most cases TAs do have control over how they present the prescribed content, anecdotal evidence and written comments at the end of the survey suggest that TAs don’t feel that they have enough ownership of the sections they teach to present material in an active manner. Adding a question about whether TAs had been instructor of record or in some way evaluate their real or perceived level of autonomy could help to clarify and further categorize survey responses. Second, we did not ask TAs what types of courses they were teaching or their duties in those courses. Most TAs teach one or more sections of laboratory courses, however some are course assistants in large lecture courses and some are instructors of record in less hands-on content-based (“lecture”-based) courses. These differences in types of courses taught could lead to very different perceptions of active learning; in the future we recommend asking in the survey what types of courses the TAs are teaching. Third, we did not specify whether or not lab activities should be considered active learning. This could lead to inflated or reduced estimates of AL among TAs based on their interpretation of AL and labs. In the future, the survey should be revised to reflect whether lab activities themselves should be considered AL. Fourth, we did not ask respondents to specify if they had participated in any teaching-related professional development activities or taken any courses focused on teaching. This information would have been helpful in determining what types of resources are being used by graduate students and how these experiences may have shaped their opinions of active learning. Adding a free-response question asking what types of training graduate students would like would help direct necessary resources to the correct Colleges and departments. Finally, asking if graduate students had taken undergraduate courses utilizing active learning would help quantify their exposure to these techniques prior to graduate school.

### Future directions

We found that STEM graduate students were simultaneously able to hold student perspectives and teacher perspectives of active learning – all that is needed is to ask them to switch roles. We also found that graduate students value active learning in their courses and are willing to teach using methods that they don’t prefer as students (although individual attitudes vary considerably). These findings spur additional questions, such as: Are the graduate students at our institution somehow different from graduate students at other universities? Recent work suggests not (46). Does acceptance of AL indicate that future faculty members will continue to use active learning in their courses if they continue to progress through academia? If these perceptions are not unique to graduate students at our institution does this mean that the future of undergraduate STEM education practices is bright? Or will our graduate students revert to more instructor-centered teaching practices that may be considered “safer” if promotion and tenure are on the line? We are unable to answer these questions with this study focused on a single time point at a single institution, but we hope that our findings are indeed indicative of a culture shift in attitudes toward teaching. We also hope that our findings spur broader studies of how graduate student professional identity develops longitudinally in different disciplines and how these teaching practices and perceptions evolve throughout their careers.

## Acknowledgements

We thank R. Seales and W. Waggenspack for facilitating the dissemination of the survey in the College of Engineering, reading previous versions of this manuscript, and many fruitful discussions of the differences among academics in various STEM fields. We are also grateful to E. Schussler and M. Chen for providing valuable comments and suggestions to the manuscript. Many thanks to the Science Education Journal Club at our institution and the S. Cotner lab for their input on previous versions of the figures and manuscript. Finally, thanks to the graduate students from both colleges who responded to the survey, without whom we would have no data.

## Supporting Information

S1 Survey Instrument

